# Sensing Traction Force on Matrix Induces Cell-Cell Distant Mechanical Communications for Self-assembly

**DOI:** 10.1101/866228

**Authors:** Mingxing Ouyang, Zhili Qian, Bing Bu, Yang Jin, Jiajia Wang, Lei Liu, Yan Pan, Linhong Deng

## Abstract

The long-range biomechanical force propagating across large scale may reserve the capability to trigger coordinative responses within cell population such as during angiogenesis, epithelial tubulogenesis, and cancer metastasis. How cells communicate in a distant manner within the group for self-assembly remains largely unknown. Here we found that airway smooth muscle cells (ASMCs) rapidly self-assembled into well-constructed network on 3D Matrigel containing type I collagen (COL), which relied on long-range biomechanical force across the matrix to direct cell-cell distant interactions. Similar results happened by HUVEC cells to mimic angiogenesis. Interestingly, single ASMCs initiated multiple extended protrusions precisely pointing to neighboring cells in distance, depending on traction force sensing. Separate ASMCs sensed each other to move directionally on both non-fibrous Matrigel and more efficiently when containing fibrous COL, but lost mutual sensing on fixed gel or coated glass due to no long-range force transmission. Beads tracking assay demonstrated distant transmission of traction force, and finite element method modeling confirmed the consistency between maximum strain distribution on matrix and cell directional movements in experiments. Furthermore, ASMCs recruited COL from the hydrogel to build fibrous network to mechanically stabilize cell network. Our results revealed for the first time that cells can sense traction force transmitted through the matrix to initiate cell-cell distant mechanical communications, resulting in cell directional migration and coordinative self-assembly with active matrix remodeling. As an interesting phenomenon, cells sound able to ‘make phone call’ via long-range biomechanics, which implicates physiological importance such as for tissue pattern formation.

## Introduction

Biomechanics and Mechanobiology has been a highly-recognized field in studying the importance of physical factors in physiologies and pathologies. Beside these extensive and elegant studies focusing on the fine local biomechanics, the long-range biomechanical force which can propagate through a relatively large scale (in hundreds of micrometers and beyond) to trigger biological responses has been an intriguing topic in recent years[1-3]. For examples, the nascent skin from the chicken embryo bears the compression force from dermal cells, which activates the follicle gene expression program[4]; cardiac cells across a wide field can synchronize their beating through mechanical communications[5]. In 1981, Harris et al. reported the morphogenesis of type I collagen (COL) fiber between two separated fibroblast explants through the cell traction[6]. In two recent reports including our own study, acini cultured from human mammary MCF-10A cells induced COL line assembly through long-range traction forces, which promoted epithelial branching pattern morphogenesis, or invasive cell phenotype[1, 2]. *In vitro* 3D culture of extracted mouse mammary glands also generated long COL fibers via cell traction forces[7]. In the past years, there have also been studies combining with computational simulations and experimental evidence that attempted to explain the mechanism of COL line formation induced by biomechanics[8-12]. So far, it still remains largely unknown how cells interact mechanically in a distant manner.

As one supporting mechanism for cell distant interactions, the locally enhanced stiffness via the nonlinear biomechanics of the newly formed COL lines could lead to directional cell migrations or cancer metastasis[2, 13, 14]. Previous work indicated that fibrillar gel could be stiffened by cell traction forces to result in cell-cell communication[11, 15]. Endothelial cell pairs showed mechanical communication on soft but not stiff polyacrylamide gel[16]. A recent study reported that cancer-associated fibroblast can align fibronectin matrix to promote directional migration of cancer cells[17]. The traction force from multiple-cell chain was also found able to add up through cell-cell junctional connections on polyacrylamide gel[18]. At current stage, the long-range biomechanical forces and the resulted mechanobiological responses still highly demand for characterization and definition. One fundamental question whether cells can directly sense the long-range biomechanical force in the matrix to result in mutual interactions and group assembly hasn’t been well answered.

Collective behavior of cell population has been well shown to exhibit during embryonic development, cell migration, immune responses, and cancer metastasis[19, 20]. Some these well-coordinated functions rely on cell-cell communications with or without direct cell contacts[1, 5, 19, 21, 22]. Extracellular matrix (ECM) is often an essential requirement for biomechanical force transmission[6, 23], which lays down a possibility to direct cell collective behaviors. Cells can recruit synthetic fibers and probe mechanics in fibrillar matrices[24]. Basement membrane (BM), which major components include collagen IV, laminin, entactin and perlecan, is abundant in epithelium, endothelium, and mesothelium[25]. Matrigel derived from mouse sarcoma cells is used as a BM mimic for *in vitro* studies, mainly consisting of two components: collagen IV and laminin[26]. Due to its structural feature, collagen IV assembles into a layer of porous reticular network as sheet-like structure with laminin[27, 28], hence less efficient in directional force transmission. COL often exists as long bundles in tissues and is able to assemble into long fibers to facilitate the efficient transmission of biomechanical force[2, 29]. Therefore, different types of ECM scaffolds such as BM and COL may have different roles in physiological events due to their biomechanical properties[30, 31]. Cells can also recruit synthetic fibers, and probe mechanics in fibrillar matrices[24]. Hence, it is an interesting topic to investigate how different ECM mechanical property, ECM remodling, and cell-ECM interaction influence long-range biomechanics and the resulted mechano-responses during cell collective behaviors.

Airway smooth muscle (ASM) cells resident in bronchial airway tissue and expressing smooth muscle alpha-actin show relatively stronger intracellular contraction force in comparison to many other types of cells in the body[32]. Although the ASM bundles are crucial in providing the needed mechanical support and contraction force for the airways in respiratory systems, excessive ASM mass is associated with airway hyper-responsiveness (AHR) in asthma attacks, due to enhanced contraction force and impaired relaxation of the airways[33]. Further study of the biomechanical property of ASMCs, especially at the population level, can have important implications in understanding the dynamic regulation of ASM structure and function, and related respiratory physiological and pathological processes.

In this work, we investigated the self-assembly behavior of ASMCs on Matrigel with or without COL. ASMCs could rapidly assemble into a well-constructed network on Matrigel with COL, which required the collective coordination of thousands of cells within 12-16 hours. This process relied on the transmission of long-range mechanical forces across the substrate to direct cell-cell distant interactions. Similar results were observed with human umbilical vein endothelial cells (HUVEC) in mimicking angiogenesis. Separated single ASMCs could sense each other in distance and initiate multiple protrusions precisely pointing to neighboring cells on both non-fibrous 3D Matrigel (without COL) and more efficiently on fibrous 3D Matrigel (with COL). Beads tracking assay demonstrated distant transmission of traction force, and mathematical modeling confirmed the consistency between maximum strain distribution on matrix and cell directional protrusions and movements in experiments. By recruiting COL from the hydrogel, ASMCs built COL fibrous network to mechanically stabilize the assembled cell network. The discovery that cell sensing of long-range biomechanics to result in precise cell-cell connections and coordinative group assembly may implicate importance in tissue pattern formation. The cell network assembly is also a useful experimental model for studying long-range biomechanics-induced mechanobiology.

## Results

### COL-mediated rapid self-assembly of ASMCs

Cells can exert collective biological functions at the population level, which can hardly be fully understood from single cell-level studies. In this work, primary rat ASMCs were seeded on 3D Matrigel containing 0.5 mg/ml COL in the circular area with 600 µm × 0.6 cm in depth × diameter (Fig. 1A). The group of cells (∼1,000-3,000 cells) rapidly self-assembled into a well-constructed network within 12-16 hours (Fig. 1B). This single continuous network incorporated nearly all cells seeded on the substrate surface, and was largely based on single cell to single cell connections. In comparison to Matrigel without COL, cells formed isolated small cysts (Fig. 1C), which indicates a necessary role for COL mediation in the assembly process.

**Figure 1.**
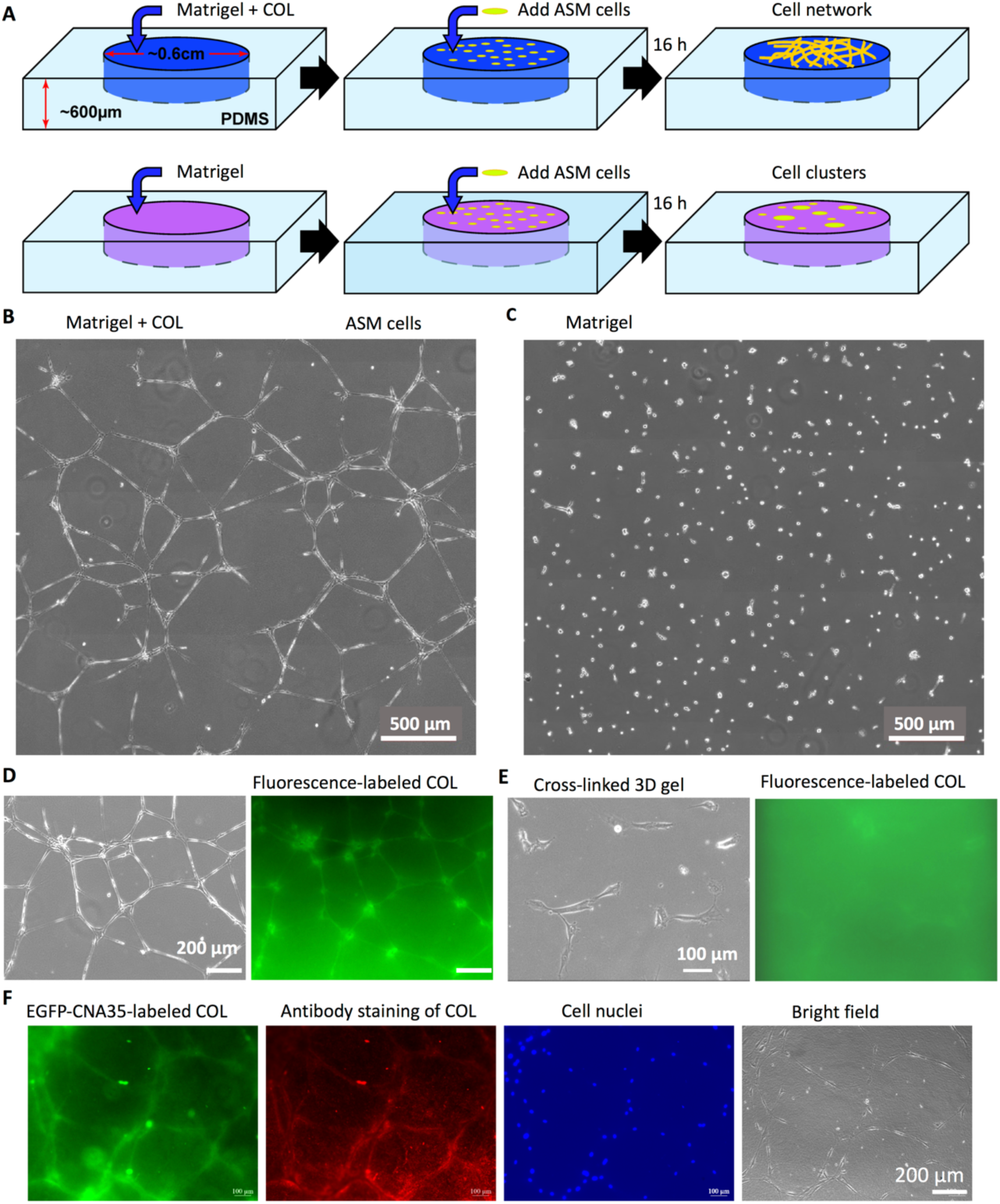
Self-assembly of ASMCs into continuous network on Matrigel substrate containing COL. **(A)** A schematic diagram depicting the experimental design with detailed steps described in the Methods. **(B)** The network morphology of ASMCs cultured on 3D Matrigel containing 0.5 mg/ml COL for 12-16 h. **(C)** The clusters morphology of ASMCs after cultured on 3D Matrigel without COL for 12-16 h. The images of (B&C) were each stitched from 9 pictures taken under 20x objective. **(D)** The nearly identical network of EGFP-CNA35-labeled COL formed along with the self-assembled network of ASMCs on hydrogel. **(E)** The morphology of ASMCs and fluorescence distribution of EGFP-CNA35-labeled COL cultured on hydrogel pre-cross-linked with 0.5% (v/v) L-glutaraldehyde for 10 min. hydrogel refers to 3D Matrigel containing 0.5 mg/ml COL in this work. **(F)** COL immunostaining by COL antibody (red) to confirm the COL fibrous network observed by EGFP-CNA35 staining (green). Cell nuclei were stained with DAPI.

To further look at the role of COL in mediating cell network assembly, COL was fluorescently labeled with its binding protein CNA35 conjugated with EGFP, which has been a well-established method[2, 34]. When ASMCs assembled into a network on top of the hydrogel, the fluorescently labeled COL displayed a nearly identical network consisting of concentrated COL lines along with the networked cells (Fig. 1D). When the proteins in the hydrogel were pre-cross-linked by L-glutaraldehyde fixation, the cells didn’t assemble into a network anymore, neither did the COL lines form (Fig. 1E). These results together suggest that cells recruited COL from their surrounding areas of the hydrogel, and COL played an active role in the cell network formation.

When cells were seeded on the hydrogel in higher density, the two nearly identical networks were formed by cells and COL (Fig. S1A&B), indicating a high capability of the network assembly. In addition, the COL fibrous network observed by EGFP-CNA35 (purified protein was displayed on Fig. S1C) labeling was also confirmed by double staining with COL antibody (Fig. 1F). These suggest that the cell & COL network morphogenesis is a robust process under the experimental condition.

### Long-range biomechanics required for self-assembly of the cell group

We then looked at the dynamic steps how the individual ASMCs assembled into a well-constructed network on the hydrogel. Through time-lapse imaging (sample size = 10), one general type of assembly is displayed in Fig. 2A: after cells were seeded on the gel, individual cells located nearby could sense each other’s locations, initiated directed extension and movement, and then assembled into one curved chain, followed by further straightening up. This process could be completed within 6 hours. Another general type of assembly occurred through cell-cell distant attractions, as demonstrated in Fig. 2B: after the first 6 hours of cell seeding, the cells already formed individual lines or local networks; then the cells with protrusive free ends could sense each other’s positions in a distant manner, and started to bend the cell bodies toward each other, and moved directionally to get connected, resulting in formation of new cell chains that were further straightened up. The full-time course of network assembling as depicted on Fig. 2A&B is shown in Movie S1, and parallel sample images of additional experiments are also shown in Fig. S2A&B. In this way of two consecutive dynamic steps, individual lines or local branching further assembled into one network incorporating nearly all cells on the hydrogel. Inhibition of cellular contraction force largely disrupted the cell alignments on the assembled network (Fig. 2C), indicating the requirement for traction force to maintain the structure.

**Figure 2.**
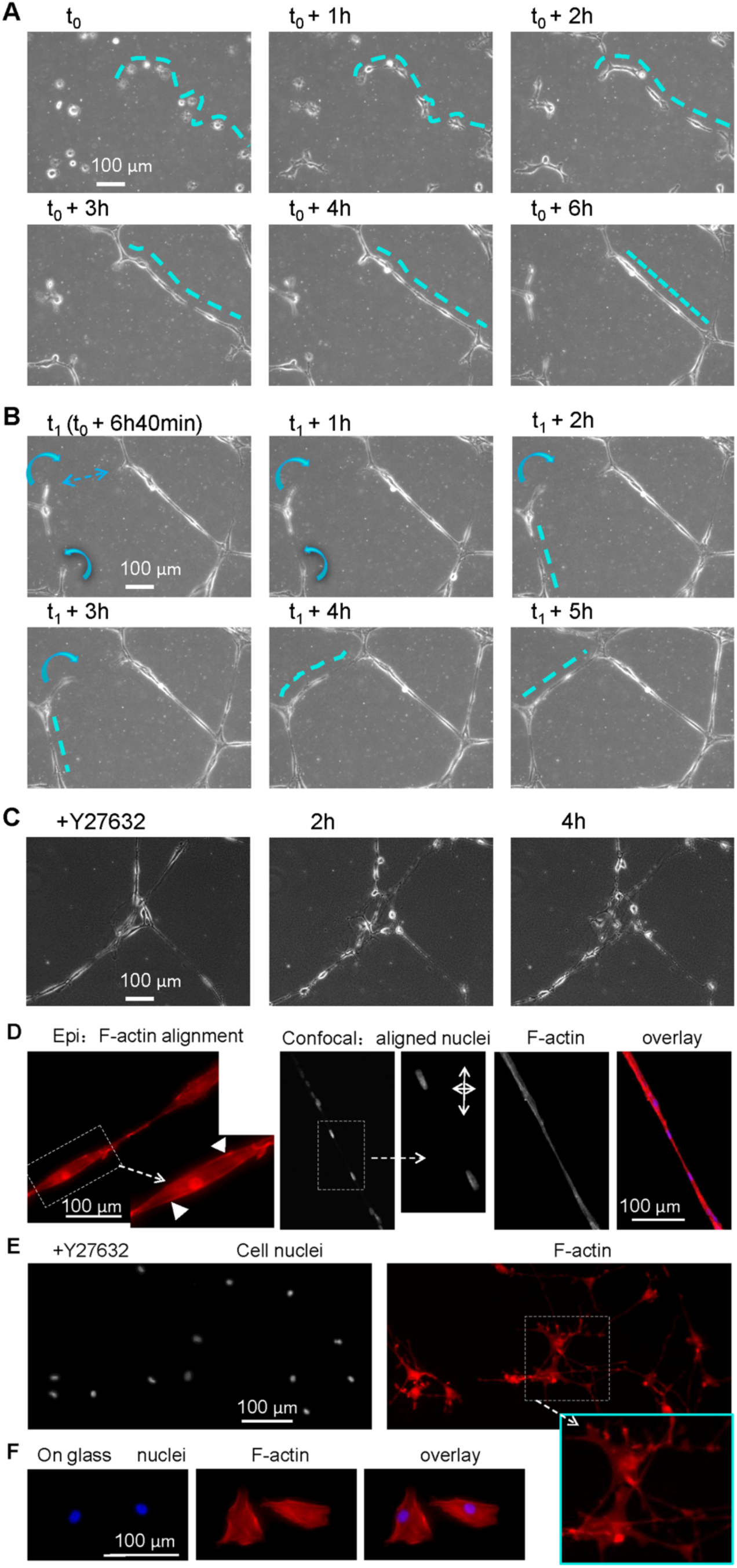
Distant interactions of ASMCs during network assembly. **(A)** The straightening alignment of a group of cells located nearby from the starting time point t_0_ to t_0_ + 6h. The broken lines indicate how the group of cells nearby got connected, formed a curved chain, and further got straightened up. **(B)** The distant attractions of cells during the network assembly. The arrows indicate that cells could sense each other distantly, followed by getting connected and straightened up. The curved arrows indicate the cell body bending to another cell. The cells on (A&B) were from the same imaging window. **(C)** Inhibition of cellular contraction force disrupted cell morphology in the network. 40 µM Y27632 was added after the cell network was assembled. **(D)** The Phalloidin staining of aligned F-actin and elongated nuclei in the well-assembled ASMCs network (cultured for over 24 h) taken by epi-fluorescence or confocal microscopy. The filled arrows indicate aligned F-actin fibers. The two crossed arrows mimic the relative length ratio along the long and short axis of the elongated nuclei. **(E&F)** The morphology of cell nuclei and F-actin staining in ASMCs cultured on hydrogel with 40 µM Y27632 (E) or cultured on regular cover glass (F).

It seems apparent that the assembly process required force involvements, due to cell activities such as the initial cell mutual sensing in distance, cell directional moving, dramatic cell body bending toward each other, and the straightening up of the cell chains. As supporting evidence to that, the ß-actin stress fibers bearing intracellular contraction were well-aligned along the direction of the cell chains on the well-assembled network (Fig. 2D), which suggests possible add-ups of traction forces. At the same time, cell nuclei were also reshaped into elongated morphology along the cell chains (Fig. 2D). When intracellular contraction force was inhibited by ROCK signaling inhibitor Y27632, neither the alignment of actin stress fibers nor cell nuclei elongation occurred (Fig. 2E). As another comparison, cells cultured on regular glass showed non-directional positioning of the stress fibers and nearly round shape of nuclei (Fig. 2F). These observations together indicate that long-range force originated from cell contraction force is required for the network assembly, and transforms the morphologies of both the cell bodies and nuclei into aligned shapes along the cell chains.

HUVECs were also applied in this system to mimic angiogenesis. On the hydrogel seeded with different densities of HUVECs, the cells were gradually assembled into thick bundles at higher density and branching at lower density in ∼30 hours, and the cell position tracking showed clearly the directional movements of cells toward the assembled structures (Fig. 3A and Movie S2). In contrast when the cellular contraction force was inhibited by addition of 40 µM Y27632, HUVECs lost persistent directional movements or branching assembly on the hydrogel (Fig. 3B), indicating similarly a requirement for cell-cell mechanical interactions. Although HUVECs didn’t form fine network-like or assemble as rapidly as ASMCs, which might be due to the different properties from cell types, these observations demonstrated the similar process to the ASMCs network assembly (shown on Figs. 2A&B and S2). It is worthy to note the cell-cell distant interactions, and the persistent directional migrations and assembly of HUVECs during the tens of hours on the hydrogel, which may implicate a mechanical principle during angiogenesis *in vivo*. For convenience, all cells refer to ASMCs in this article unless notified.

**Figure 3.**
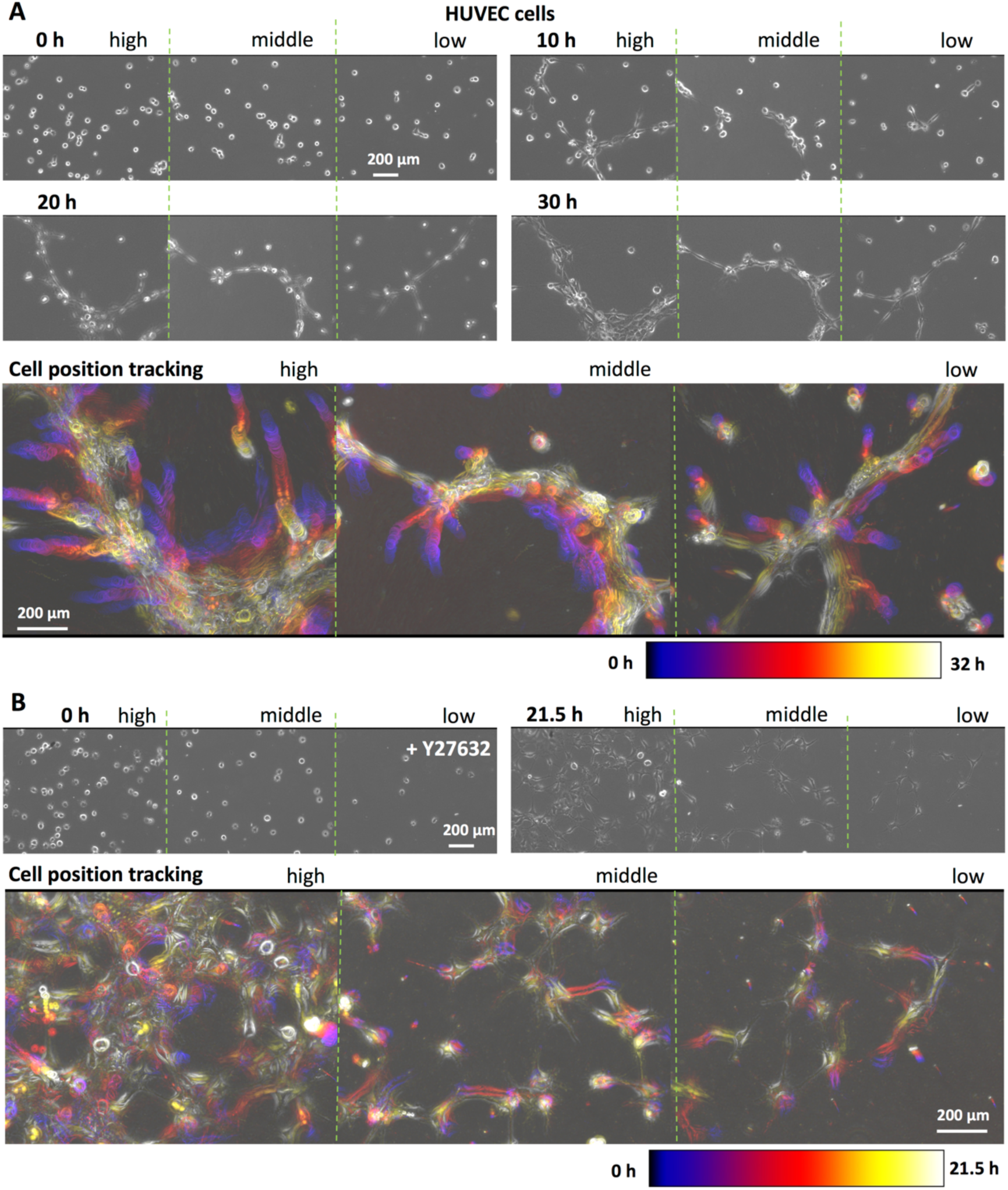
Traction force-regulated directional movements and assembly of HUVECs on the hydrogel. HUVECs were seeded at different densities (high, middle, low as indicated) on 3D Matrigel containing 0.5 mg/ml COL. **(A)** Time-lapse imaging (interval = 30 min) revealed the dynamics processes of cell-cell distant attractions, cell directional movements, and branching assembly. The cell position tracking was done by the function ‘Time-lapse color coder’ in the software ImageJ, which showed the cells’ persistent directional migrations toward the assembled structures during the 32-hour period. **(B)** The losing cell directional movements, or branching assembly of HUVECs on the hydrogel with the addition of cellular contraction force inhibitor Y27632 (40 µM) into the culture system. The cell position tracking was done during the 21.5-hour period.

### Traction force-dependent cell directional protrusions toward neighboring cells

To examine the process how ASMCs originally sense each other’s positions in distance, we seeded cells at low density (∼3,000 cells/cm^2^) on the COL-containing 3D Matrigel, which could also allow longer duration for cell mutual interactions. Surprisingly, it appeared that a single cell was able to initiate multiple extended protrusions precisely pointing to neighboring cells, as one representative sample shown in Fig. 4A and Movie S3. In this case, one single cell started 3 directional protrusions within 2 hours upon seeding on the hydrogel, which continuously extended toward three other cells in the following hours; another cell in the same field was able to initiate 4 protrusions toward four different cells in distance after 13 hours from the seeding time, and got connected with these cells in the next ∼6 hours; eventually, all these cells in the field were able to get connected together within ∼20 hours from the seeding time.

**Figure 4.**
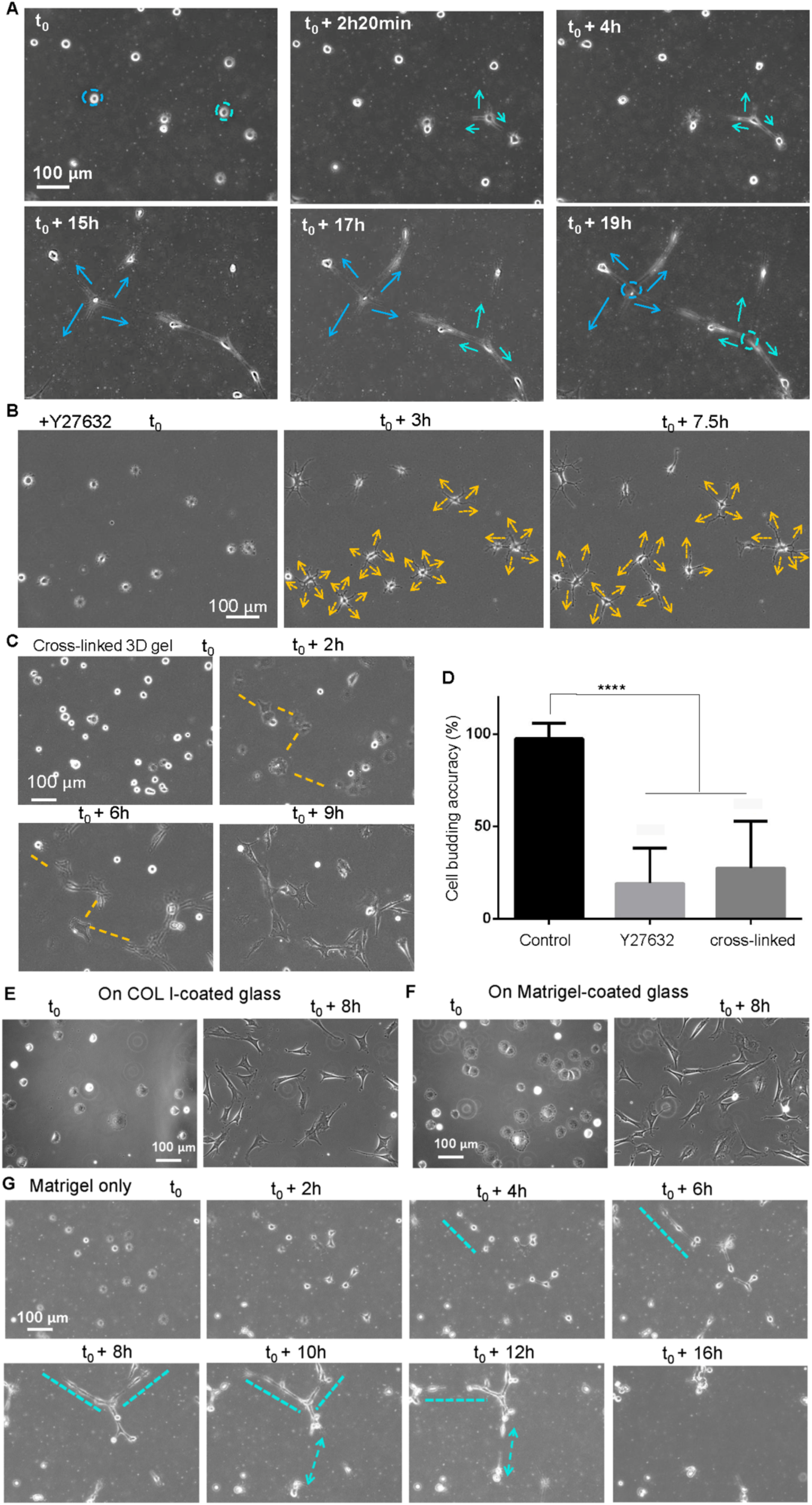
Traction force-dependent cell directional protrusions and movements toward neighboring cells. **(A)** The directional protrusions of ASMCs on hydrogel pointing to neighboring cells in distance during network assembly. The arrows indicate the multiple protrusions from these two leading cells pointing to neighboring cells in distance. **(B)** The random protrusions of cells on the hydrogel with addition of contraction force inhibitor Y27632 (40 µM). The arrows indicate the protrusion directions irrelevant to the positions of other cells nearby. **(C)** The lost mutual sensing between cells after cross-linking of the hydrogel by 0.5% L-glutaraldehyde treatment for 10 min. The broken lines indicate the distance between cells located nearby. **(D)** Quantification of cell protrusion accuracy (mean ± S.D.) on hydrogel under the indicated conditions, according to whether the cell protrusions pointed to neighboring cells, either accidentally or by distant attraction. N = 100 cell samples for each condition, and **** indicates P value <10^−4^ through the Student’s t-test analysis. **(E&F)** No mutual sensing or distant attraction between cells cultured on COL-coated (E) or Matrigel-coated (F) glass. **(G)** The branching assembly through cell-cell position sensing and directional movements on non-fibrous 3D Matrigel without COL.

One interesting question is what helped the cells to sense each other’s locations by which directional protrusions occurred upon seeding on the hydrogel. We found that when cell contractility was pharmacologically inhibited by Y27632, although cells could still initiate multiple protrusions, they were unable to sense each other anymore because the protrusions lost their directions pointing to neighboring cells regardless of the cell seeding density (Figs. 4B, S3, and Movie S4). This suggests that cell traction force was necessary for cell mutual distant sensing on the hydrogel. We further tried to inhibit the force transmission efficiency across the hydrogel by pretreatment of the gel with L-glutaraldehyde. When L-glutaraldehyde fixation caused proteins cross-linking in the hydrogel, it abolished the cells’ capability to sense each other’s locations even at close positions, or assemble into a network (Figs. 4C, S4, and Movie S5). Based on quantification from 100 cell samples, nearly 100% of the sustainable protrusions initiated from single cells were accurately pointing to neighboring cells, whereas inhibition of cell traction force or force transmission in the matrix abolished the directional protruding (Fig. 4D). This data indicates that for cell-cell distant sensing, it is necessary for traction force to reach neighboring cells via the matrix deformation and strain. Since hard glass surface doesn’t transmit cell traction force to neighboring cells while not stopping mutual possible chemical or electronic signal passing between the cells, we further proved that the cells cultured on COL or Matrigel-coated glass couldn’t sense each other’s positions nearby or assemble into network (Fig. 4E&F, and Movie S6). From above experimental observations, we conclude that cell sensing traction force in the matrix is required to induce cell-cell distant communications for the network assembly.

To check whether COL was absolutely required for cell-cell distant sensing, we seeded cells on the hydrogel containing Matrigel only. Time-lapse imaging revealed that cells were still able to sense other cells’ locations nearby with directional protrusions at lower or higher seeding density, but less efficiently than on the hydrogel containing COL (Figs. 4G and S5, and Movie S7). This may be explained by the fact that the non-fibrous Matrigel is structurally inefficient in directionally transmitting cell traction force. These do indicate that traction force sensing on the matrix is essential for cell-cell distant communications with or without COL fibers in the hydrogel.

### Stabilization of the cell network by COL remodeling

Although cells were able to sense each other and get connected on the non-fibrous Matrigel without COL, they couldn’t maintain stable cell network, as the connected cell chains eventually collapsed into cell clusters (Fig. 4G). In considering that fluorescence-labeled COL formed one nearly identical network along with the one of ASMCs (Fig. 1C), COL fibers might be critical in stabilizing the cell network.

Thus, we used time-lapse imaging to simultaneously check the cell directional protruding, the assembly of ASMCs network together with the formation of fluorescent COL fibrous structure. As shown in Fig. 5A, one individual cell initiated three protrusions pointing to neighboring cells, but at this initial stage, there were no obvious concentrated COL lines formed between the mutually attracted cells. As the cells continued to extend to each other, COL lines started to appear and became increasingly visible afterward (from t_1_ + 120min to t_1_ + 240min in Fig. 5A). As shown in Fig. 4B, fluorescent COL showed accumulation in line with the cell chains during the 3 to 15 hours of cell seeding. This time sequence of COL gradual accumulation between the two mutual-interacting cells in Fig. 4A was also confirmed by the fluorescence intensity quantification (Fig. 5C). It is noticeable that COL was accumulated beside the cell chains but not overlapped with the cells, which was also confirmed by the fluorescence intensity quantification (Fig. 5D), indicating the appearance of COL lines after the cell chains. Additional sample of directional protrusions along with COL line formations can be seen in Fig. S6. It should be noted that there was certain fluorescence photobleaching during the time-lapse imaging, as EGFP was added into the gel, but not continuously secreted from the cells.

**Figure 5.**
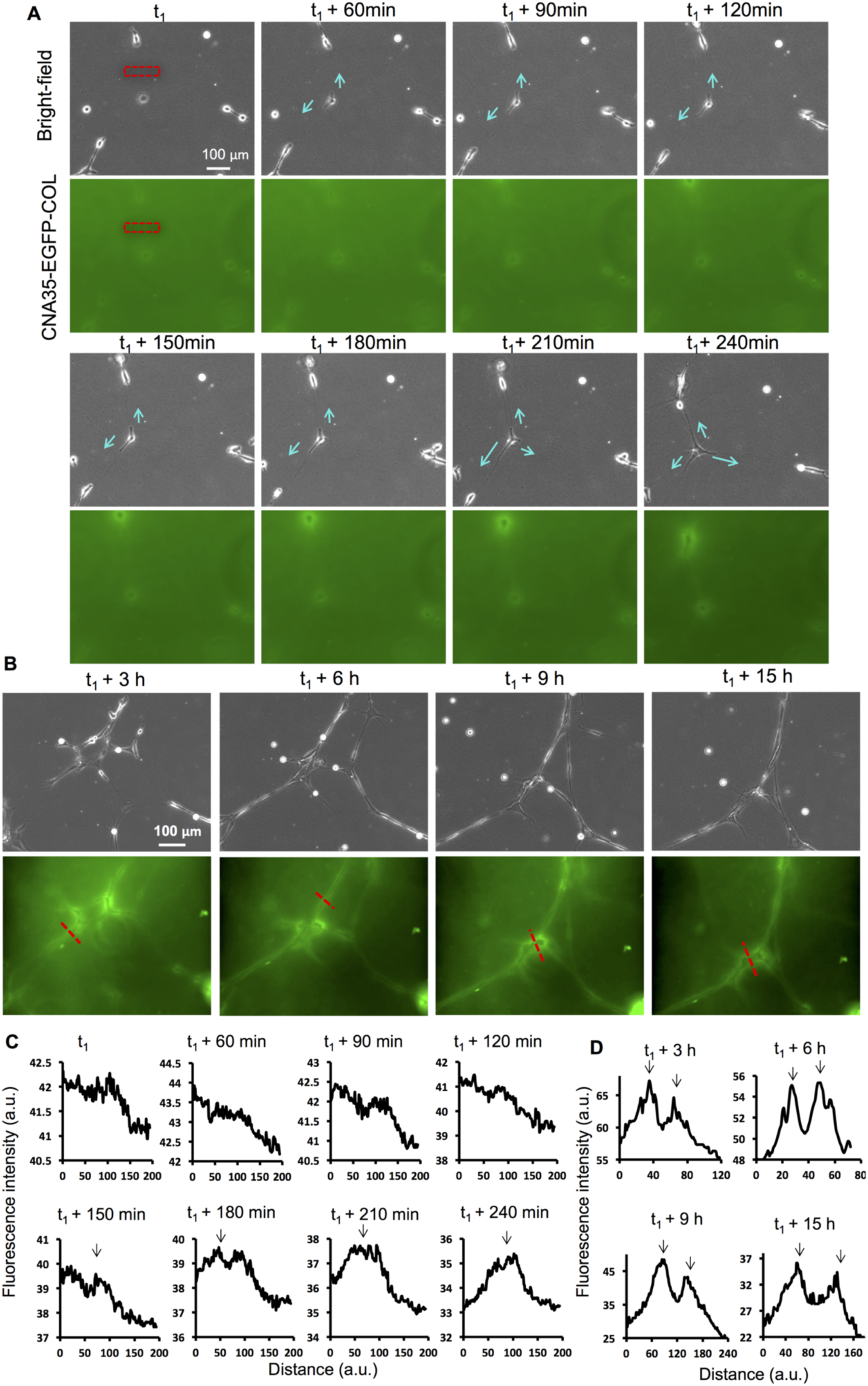
Cell directional protrusions, and subsequent emergence of COL fibers between neighboring cells. **(A)** The time-lapse imaging of the cell which initiated three buds accurately pointing to neighboring cells. The arrows indicate the directional budding. **(B)** Simultaneous imaging of the cell network assembly along with the COL network formation. **(C)** The quantifications of fluorescence intensity in the left-right direction along the selected rectangular region. The arrows indicate the gradual accumulation of COL between the two cells with distant sensing. **(D)** The quantifications of fluorescence intensity along the labeled lines across the cells. The two peaks on the curves (shown by the arrows) indicate the accumulations of COL beside the cell chains. a.u. refers to arbitrary unit.

This observation above confirmed that ASMCs recruited COL from the hydrogel to form a similar COL network in the substrate. In considering that cells couldn’t assemble stable networks on Matrigel when COL was absent or fixed by L-glutaraldehyde, the fibrous COL structure must be crucial in mediating and maintaining the cell network.

### Demonstration of long-range force transmission across the matrix

Traction force microscopy (TFM) technique has been widely used to study the cellular traction force exerting on the matrix based on measurement of local substrate deformations[35]. By applying the similar principle, we visualized the long-range force transmission across the matrix. Briefly, when fluorescently labeled beads were embedded in the hydrogel, their position movements could then be tracked during the network assembly of ASMCs. Fig. 6A shows a map of local deformation of the hydrogel based on the bead tracking analysis during a typical 5-hour period chosen from the time course in Fig. 2B. It can be seen in the square-labeled Region 1, the local cell network was nearly completed, but beads didn’t show significant deformations within the 5 hours, indicating little active local assembly due to possible balance of force in different directions. In Region 2, the cells were connected and assembled into network within the 5 hours, and the beads moved a lot toward the cells, as groups of local beads shifted their positions in the similar direction, indicating strong long-range traction force transmission in the matrix from the cells.

**Figure 6.**
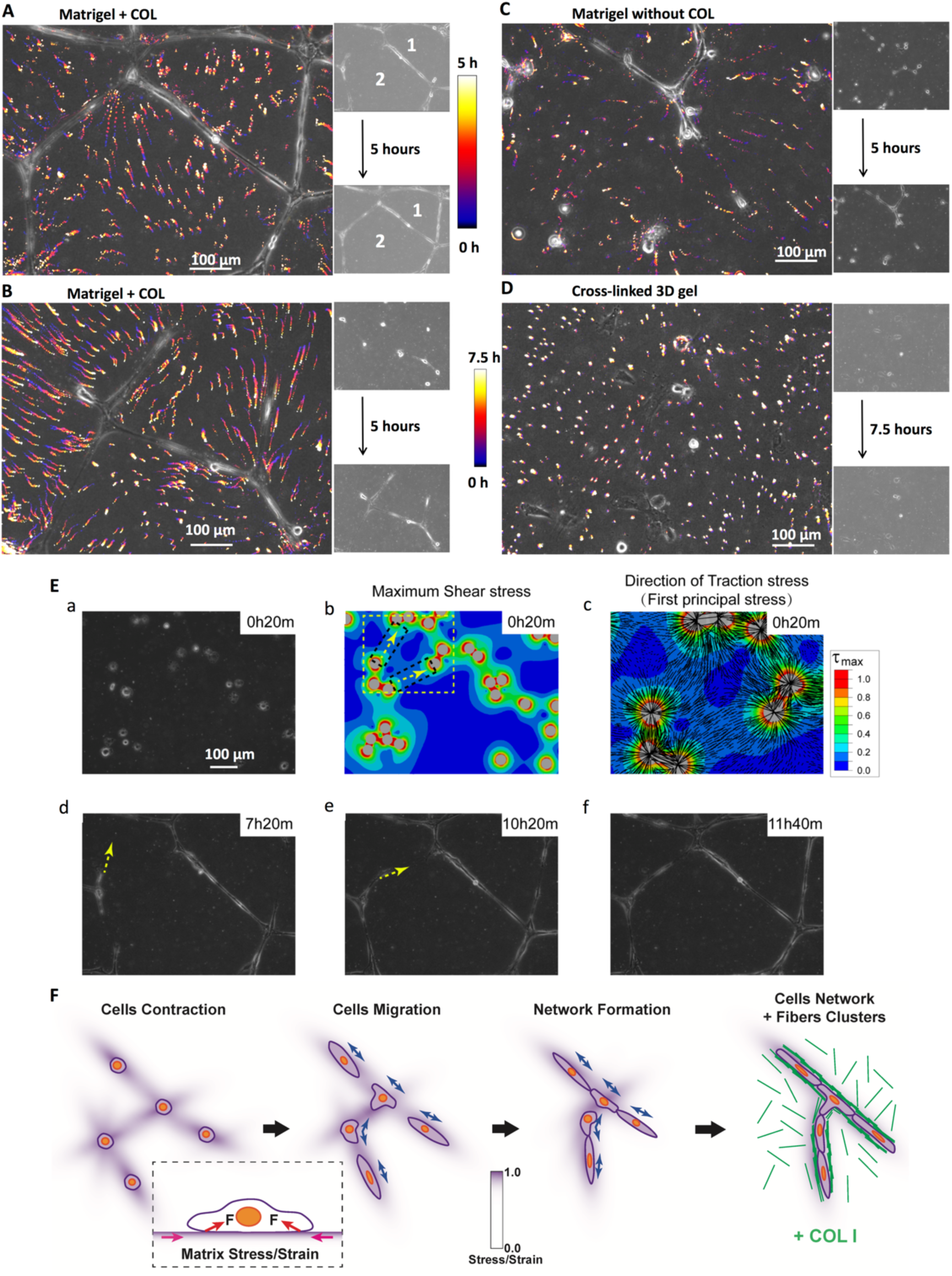
Long-range traction force transmission across the matrix, and the consistency between network assembly and maximum stress distributions from FEM modeling. **(A-D)** The positions of beads on the hydrogel were tracked along the indicated period of cell assembly, and the pseudo-color time scale represents the positions along the time points. The 5 or 7.5-hour beads tracking analysis was shown from the sample images of Figure 2B (A), Figure 3A with low cell density (B), Figure 3G on Matrigel without COL (C), or Figure S4B on cross-linked hydrogel by L-glutaraldehyde (D). Regions 1 and 2 indicate less or more bead-position shifts. **(E)** The consistency of directions in cell migration and modeled maximum stress distributions. (a, b) Individual cell positions and the maximum shear stress distribution on matrix at t_0_+20min (also displayed in Fig. 2a). (c) The directions of traction stress (first principal stress) in the yellow rectangle region of panel (b) were shown as black lines. (d) & (e) Two possible directions of cell protrusions (yellow arrows), and (f) network formed along the directions of panel (e). The pseudo-color scale bar represents increasing relative maximum shear stress at local place. **(F)** The diagram of traction strain-induced cell-cell distant communications and network assembly. Cell contraction force exerts non-uniform stress and strain distribution on matrix surface; cells mechanosense the uneven strain distribution, and initiate directional protrusions and migration, resulting in mutual connections and network assembly; cells recruit COL from the matrix to assemble fibrous structure in mechanically supporting cell network.

Fig. 6B shows the beads tracking-based deformation map of the hydrogel during a 5-hour period chosen from the time course in Fig. 4A where cells were seeded at low density (∼3,000 cells/cm^2^). The cells started largely from single cells to get connected, and there were significant movements of beads located nearby or distantly toward the cells during cell assembling, and the beads located beside the cells might move a lot along the cells. This demonstrates that the traction force from only a few cells could propagate to distant locations through the matrix. In contrast, when the hydrogel was fixed by L-glutaraldehyde cross-linking, positions of the beads stayed almost stationary without significant shifts in the 7.5-hour period as shown in Fig. 6C, which indicates that the force transmission across the matrix was inhibited in the cross-linked 3D gel, supporting no cells or COL network assembly (Figs. 4C and S4). In the case of 3D Matrigel without containing fibrous COL where cells were still able to sense each other in a distant manner as shown in Figs. 4G and S5, the bead tracking analysis also revealed distant force transmission in supporting that observation (Fig. 6D). However, it is noted that the beads shifts on Matrigel only were less dramatic than those on Matrigel containing COL (Figs. 6B vs. 6D). These data together strongly support that sensing traction force initiates cell-cell distant communication on the hydrogel substrate.

### The correlation between cell distant interactions and stress distribution in the matrix

The above results demonstrated that cell-cell distant attractions and the network assembly required cell traction and force transmission across the matrix, which was verified by a global displacement or strain field of the matrix induced by cells migration and contraction. We then took a further step to check the relationship between the cell directional protrusive extending and movement and the global stress distribution on the matrix. For the lack of solving methods about inhomogeneous nonlinear stress-strain inverse problems, a simplified finite element method (FEM) model was introduced to investigate this relationship, as similarly applied in previous work[36]. The modeling results confirmed that the direction of cells protrusion and network formation was in line with the distribution of maximum shear stress and directions of the traction stress (also known as first principal stress, which was consistent with the directions of first principal strain at local place in such condition) around the cells (Fig. 6E). Cell groups in the region with higher maximum stress distribution had a high propensity to protrusion and communication with each other, which led to the formation of network subsequently (see Fig. 6E and Fig. S7). As shown by Fig. 6E(a-d), it predicted that cells on the top-right side assembled into a long cell-chain connection, which formed in 3-6 h in experimental observation (see Fig 2A). The down-left region also had a possibility to protrude towards top-right region due to the higher maximum stress distribution and directions of first principal stress (see Fig. 6E(b-e)), which was verified by the experimental results on Fig. 2B. The cell protruded to these two directions and formed network connection with the cells on the top-right region in ∼11 h (see Fig. 2B). At low cell density condition, cell might protrude to several directions (see Fig. S7). The protrusion directions also confirmed the prediction from the modeling that the direction of cells protrusion and network assembly was related to the maximum stress distribution. The modeling results, therefore, confirmed that the cell preferred to protrude and assemble network in the regions with higher maximum shear stress and toward the directions of first principal stress.

Taken together, we propose the mechanism of the network morphogenesis of ASMCs in terms of long-range force transmission and cell traction force sensing (reflected by the distributions of both stress and strain) on the matrix, along with mechanical support from COL fibrous structure. As depicted in Fig. 6F: cell contraction force exerts non-uniform stress and strain distribution on matrix surface, and the traction strains are transmitted through the matrix to reach others cells; cells mechanosense the uneven strain distribution, and initiate directional protrusions and migration, resulting in mutual connections and network assembly; cells recruit COL by remodeling the matrix to assemble fibrous structure in mechanically supporting cell network.

## Discussion

This work demonstrated that cells are able to sense the traction force in the matrix to induce distant mechanical communications, resulting in cell directional protrusions, migrations and connections. The cells responded to traction force on the matrix for distant communications, and induced COL remodeling in the hydrogel. Fibrous COL displayed important roles in facilitating the efficiency of force transmission to induce the assembly and stabilize the cell network. These findings may help advance the understanding of the feature and function of long-range biomechanical force at the cell population level. The observed high mechano-sensitivity of ASMCs might also suggest a re-enforced feedback of enhanced contraction by excessive ASM under asthmatic condition.

### 1. Long-range biomechanical force induces cell-cell distant communications and collective behaviors of network assembly on hydrogel

During embryogenesis and body development after birth, cells are often organized into groups to function collectively. For instances, the group movement of cells in developments[37], the nerve cells’ complex signal process in the brain such as called neuronal circuits[38, 39], the synchronized beating of cardiomyocytes in the heart[5], and the targeting responses of immune cells[40]. From the past research, spatial biochemical signals have been discovered as guiding factors in coordinating the cell collective activities[41, 42]. Recent evidence has added an emphasis on biomechanical factors in coordinating the cell collective activities [4, 43, 44]. Here we focused on the characterization and mechanobiology of long-range biomechanical force which could be transmitted across a group of cells at millimeter scale[1, 2, 12, 13]. We found that long-range biomechanics induced cell-cell distant attraction and directional movement which were essential for the rapid assembly of thousands of cells on hydrogel (Figs. 1 & 2). Even at low cell-seeding density, beads tracking assay demonstrates that the cell traction force could reach hundreds of micrometers away and beyond on the matrix (Fig. 6B). Inhibition of traction force or force transmission diminished cell-cell distant sensing and the assembly process (Figs. 1D, 4B-D, S3 and S4). Similar results were also observed by HUVEC cells in mimicking angiogenesis (Fig. 2A&B). These indicate that force transmission across long distance on the matrix is required for the observed cell-cell distant communications and collective behaviors.

### 2. ASMCs sense traction force in matrix with directional protrusive extension and distant attractions

One fundamental topic in mechanobiology is how cells respond to biomechanical force. As surprisingly observed here, ASMCs could extend multiple protrusions precisely pointing to neighboring cells (Figs. 4A&D, and 5A). This indicates one way how cells react to traction-induced strain passing through the matrix. Previous studies based on experimental measurement and computational simulations also demonstrated that traction-induced strain was more concentrated on the region between cells seeding on COL gel than on other surrounding regions[2, 9]. There has also been evidence that cells would migrate in the direction with more concentrated COL fibers or higher substrate rigidity[11, 20]. In our observation here, single cells could sense each other distantly and extended directional protrusions as quickly as within 2 h seeding on the hydrogel (Fig. 4A). There was no visible or measurable COL accumulation in the direction when protrusive extension occurred (Figs. 5A & S6A). At the same time, on 3D Matrigel without fibrous COL, cells could also sense each other in distance (Fig. 4G). On hard glass coated with COL or Matrigel, although cells located nearby could still communicate with each other through possible chemical or electronic signals, but couldn’t sense each other anymore without the transmitted traction-induced strain through 3D matrix (Fig. 4E&F). Together in considering the distant transmission of single cell traction force (Fig. 6A-D), ASMCs are able to sense the dynamic traction force induced on the matrix in a quick manner, and respond with directional protrusive extension toward neighboring cells. We further confirmed that the protrusive extension process relied on cell traction force and force transmission in the matrix (Figs. 4B&C, S3 and S4). FEM modeling confirmed the correlation of maximum traction force-induced in matrix and the directions of cell protrusions and movements (Figs. 6A-D & S7). The mechanism of directional protrusive extension in response to traction force needs further investigation.

### 3. ASMCs actively recruit COL from the hydrogel to assemble COL fibrous structure which provides mechanical support in maintaining the cell network

Collagen bundles as the most abundant collective tissue component provide mechanical property and structural supports for tissues *in vivo*[45]. In our simple *in vitro* culture system, when ASMCs were self-assembled into a well-constructed network on hydrogel, a nearly identical COL network was also built along the cell network (Figs. 1C, 5B & S1B). This demonstrates that cells can recruit COL from the hydrogel to assemble COL fibrous structure, indicating an active interaction between cells and ECM, and a possible model for *in vivo* condition. When the hydrogel didn’t contain COL component, ASMCs failed to maintain an assembled network, instead collapsed into clusters, although cell-cell distant attractions still occurred at certain level (Figs. 4G and S5). Hence the assembled COL fibrous structure provides mechanical support in maintaining the cell network.

### 4. The implication from the mechano-property of ASMCs as ‘re-enforced feedback’ in chronical asthmatic condition

Asthma is a common disorder of the respiratory system, affecting about 300 million people world widely with increasing prevalence[46]. The most critical asthmatic symptom is airway hyper-responsiveness (AHR) which is largely derived from enhanced ASM mass and prolonged ASM contraction[33], hence the mechano-property of ASM is relevant to asthma treatment. Our work here shows that ASMCs displayed both strong contraction force and high mechanosensitivity. The cells responded to traction force transmitted distantly in the matrix with directional protrusions, movements and cell shape change, which further led to cell-cell connections and straightening-up of the multiple-cell chain (Figs. 2A&B, and S2). This mechanical property of ASMCs suggests that the appropriate amount of ASM in airways sounds critical in maintaining the normal function of bronchial tubes. In other words, enhanced ASM might lead to a possible ‘re-enforced feedback reaction’ in which enhanced traction force in airways would further activate the mechano-signals of ASMCs, and induce prolonged contraction responses leading to AHR.

Together, this work has further characterized the long-range biomechanical force in coordinating cell collective behavior and inducing mechanobiological response in a distant manner, which fundamental principle may shed new light on understanding tubulogenesis, neuronal network morphogenesis, and cancer metastasis. Interesting insights have been provided on the active interactions between cells and ECM components, and in understanding the AHR symptom in asthma.

Whether cells can directly sense force from their microenvironment has been an interesting long-standing topic[47]. We believe that this work has helped generation of new knowledge in revealing that cells are capable of sensing traction force in matrix to induce cell-cell distant mechanical communications, which may be critical for tissue pattern formation in vivo. Hence, this lays down another important principle beside cell rigidity sensing for cell distant mechanical interactions in the field of biomechanics and mechanobiology. Further prospective research is expected on exploring the intracellular mechanism of the distant biomechanical force sensing. How the fundamental principles of long-range biomechanics occur *in vivo* also waits for investigations in more complex system.

## Materials and Methods

### Cell culture and reagents

Primary airway smooth muscle cells were originated from 6-8-week old female Sprague Dawley rats, as described before[48]. The cells were cultured in low-glucose DMEM (Invitrogen) supplemented with 10% FBS, and penicillin/streptomycin antibiotics. ASMCs applied in the experiments were generally within 10 times of passages during regular culture. Y27632 was purchased from Santa Cruz Biotechnology; mouse monoclonal anti-collagen type I antibody (Clone COL-1), Rhodamine-conjugated goat anti-mouse IgG Antibody, Phalloidin and DAPI were from Sigma; red fluorescent beads (FluoSpheres™ Polystyrene Microspheres, 1 µm in diameter) were from Invitrogen.

### Preparation of hydrogel in polydimethylsiloxane (PDMS) mold and the culture of cell networks

A thin layer of PDMS (∼600 µm in thickness) was generated by addition of 5 ml of the two well-mixed liquid components from the Sylgard 184 kit (Dow Corning) (at 10:1 mass ratio) onto a 9 cm-diameter dish, followed by curing at 70°C for 3-4 h. The PDMS sheet was cut into circular pieces in order to fit onto the glass-bottom dishes with 2.0 cm glass surface in diameter (NEST). One or more holes with 0.6 cm in diameter were created on those pieces by a mechanical puncher. Then the PDMS was sterilized by 5-min solicitation in 75% ethanol and exposure to UV light for hours or longer in the cell culture hood. The mold was assembled by attaching the PDMS piece onto the center of glass-bottom dish.

To prepare the hydrogel, about 20 µl of Matrigel solution or Matrigel containing 0.5 mg/ml COL (mixed at 1:1 volume ratio) was added into the PDMS mold on ice, which got gelled in the 37°C incubator for 30 min. To seed cells, about 30 µl of cell suspension was added on top of the hydrogel, and stayed for 10 min in the incubator before adding more culture medium.

For cross-linking, the pre-made hydrogel was incubated in PBS solution containing 0.5% (v/v) L-glutaraldehyde for 10 or 30 min, followed in the order by three times of washes (5 min each) with PBS solution, incubation in 50 mM Tris-HCl (PH 7.4) for 1 h to neutralize the left glutaraldehyde, and another three times of washes (5 min each) with PBS solution.

### Producing EGFP-CNA35 protein from *E. Coli*, and COL labeling

The pET28a-EGFP-CNA35 DNA construct purchased from Addgene was described before[34]. The procedures for expression and purification of the EGFP-CNA35 protein has been detailed in our previous publications[49]. In brief, the DNA construct was transformed into BL21(DE3) competent *E. Coli*, and the protein expression was induced by Isopropyl β-D-1-thiogalactopyranoside (IPTG) in liquid culture at room temperature (∼25°C). After lysis of the bacterial pellets in B-Per protein extraction reagents (Thermo), the 6xHis-tagged EGFP-CNA35 protein was pulled down by HisPur Ni-NTA agarose resin (Thermo), and purified by nickel chelation chromatography. The purified protein solution was examined by 10% SDS-PAGES (sodium dodecyl sulfate-polyacrylamide gel electrophoresis) and Coomassie Blue staining, which showed only one major band with molecular size of ∼60 kD (Fig. S1C). The protein concentration was determined by Bradford Protein Assay Kit (Thermo).

To label COL, purified EGFP-CNA35 protein was incubated with COL solution at a mass ratio of 1:20 on ice for 10-15 min, followed by neutralization with a basic solution and further mixing with Matrigel solution for gel formation at 37°C.

### Epi-microscopy for live cell imaging, and confocal imaging

The live-cell epi-microscopy system (Zeiss) was equipped with X-Y-Z stage for multiple-position imaging, fine auto-focusing function for long duration of time-lapse imaging, and temperature-CO2 control chamber to maintain cell culture condition. The stitching function was applied to capture the entire cell network growing on the hydrogel. Most of the imaging experiments on hydrogel were conducted under a 20x objective. During the time-lapse fluorescence imaging of COL, the excitation light from the lamp was reduced to 1/16 of the full power, and interval time was set as 30 min in order to minimize the photo-bleaching issue.

Confocal microcopy purchased from Zeiss was applied to scan DAPI-stained cell nuclei and Phalloidin-stained actin fibers in the experiments.

### Beads tracking assay during the cell network assembly

Red fluorescent beads (1 µm in diameter) were mixed into the hydrogel at ∼1/180 volume ratio, and the beads were visible under the bright field of the microscopic images. In our experiments during the 12-16 h time-lapse imaging, we tracked those bead positions located on the surface of the hydrogel and visible under bright field, which could avoid the toxicity from exciting the fluorescent beads during the long duration.

For position tracking analysis, the beads on the bright-field images were shown up obviously through contrast adjustments, followed by binary processing in the software Photoshop. Then the beads on each image were assigned with pseudo-colors from cold to hot following the 5h or 7.5h time sequence order. Afterwards, individual images of the time sequence were overlaid and merged into one picture to display the bead movements along the duration time. Individual images could also be assembled into movie file to show the dynamics of bead positions.

### Modeling of traction-induced strain distribution on matrix with FEM

A simplified FEM model was introduced to investigate the correlation between matrix stress distribution and the direction of cell protrusion and network formation. The interaction surface of cells and matrix was reduced to an isotropic linear elastic plane surface. Cells region was removed and traction force was applied on the edges of cells to mimic the contraction force of cells on the matrix surface. The FEM model was solved by ABAQUS packages. The stress value was normalized to reach a relative quantitative analysis and results.

### Data quantification and statistical analysis

The fluorescence intensity of labeled COL was measured by the function of Plot Profile in ImageJ. The cell protrusion accuracy was calculated based on 100 cells which extended protrusions when seeding on the hydrogel. Those protrusions pointing to neighboring cells were counted as directional protrusions, either accidentally or due to distant attractions. Student’s t-test was applied for statistical analysis, and mean ± S.D. (standard derivation) was presented. P value < 0.05 indicated a significant difference in statistics during the comparison.

Additional materials and methods are available in the online supporting information.

## Acknowledgements

we appreciate Drs. Xiang Wang, Mingzhi Luo, Che Zhao and Jia Guo, and Jingjing Li, Jin Dong (Changzhou University), and Yinxiu Chi (Central South University) for technical supports and other assistance. This work was supported financially by Natural Science Foundation of China (NSFC 11532003, 11872129, 31670950), Natural Science Foundation of Jiangsu Province (BK20181416), and Changzhou Science and Technology Bureau (CZSTB CZ20180017).

## Author contributions

M. Ouyang and L. Deng conceived the project, and designed the research; Z. Qian performed the major experiments and partial data analysis; M. Ouyang did the major data analysis & organization; B. Bu conducted FEM modeling of stress distribution with written description; Y. Jin conducted bead tracking assays; J. Wang performed COL staining and HUVECs experiments; L. Liu assisted in setting up equipment; Y. Pan assisted in confocal imaging and atomic force microscopy; L. Deng provided major financial support and the setups of equipment; M. Ouyang and L. Deng wrote the paper.

