## Supplemental figures and movie legends for "Sensing Traction Force on Matrix Induces Cell-Cell Distant Mechanical Communications for Self-assembly"

Supplementary Materials including:

Supplementary Figures S1-S7;

Supplementary Movies Legends S1-S7.

### Supplementary Figures

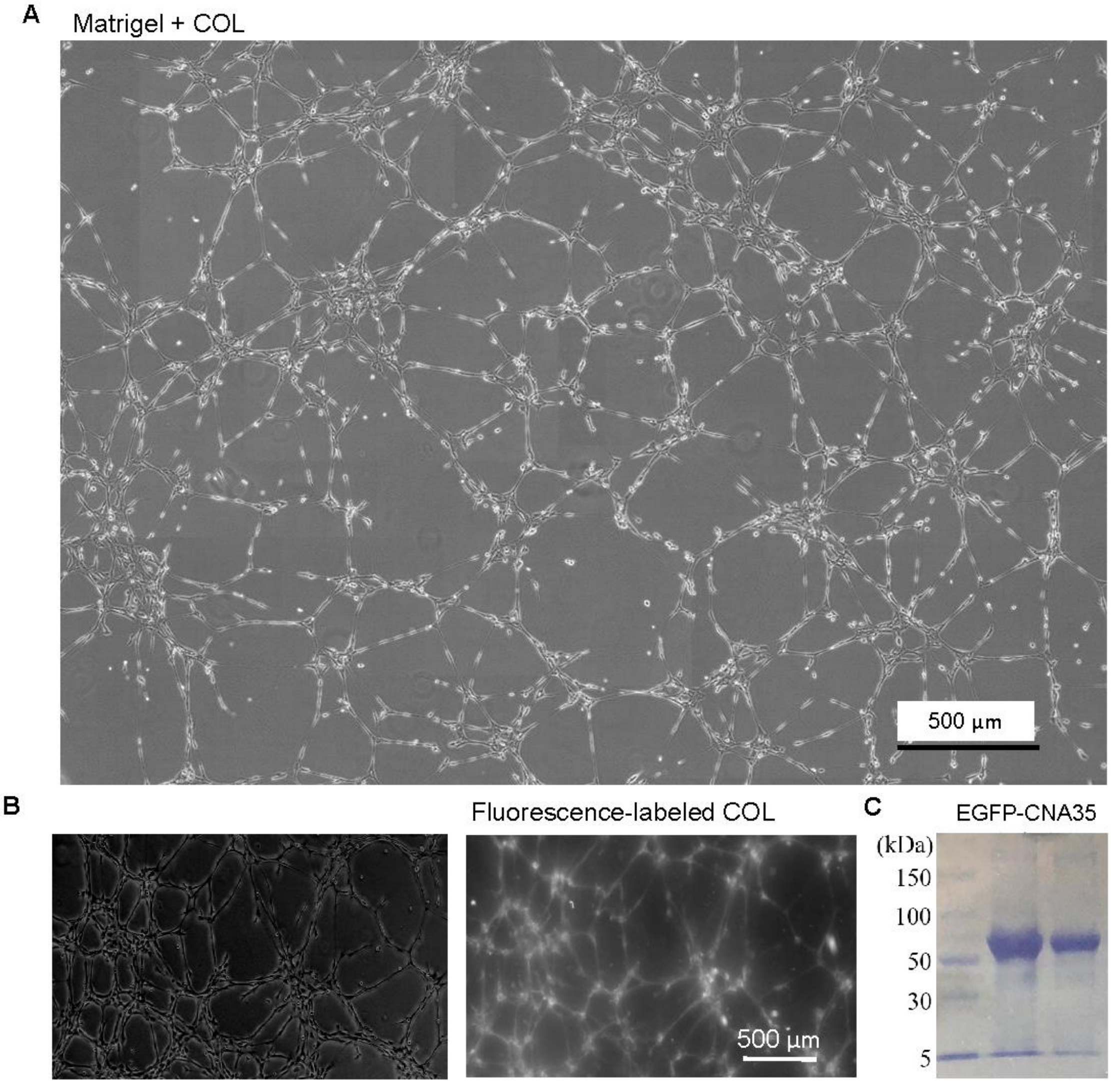


**Figure S1. The morphologies of ASMCs network and COL fibrous network. (A)** The assembled network of ASMCs at higher cell-seeding density (~9,000 cells/cm2). **(B)** The COL fibrous network along with the cell one under higher cell density. COL was labeled with EGFP-CNA35 in 3D hydrogel. **(C)** The purified EGFP-CNA35 protein resolved on 10% SDS-PAGE with Coomassie Blue staining.

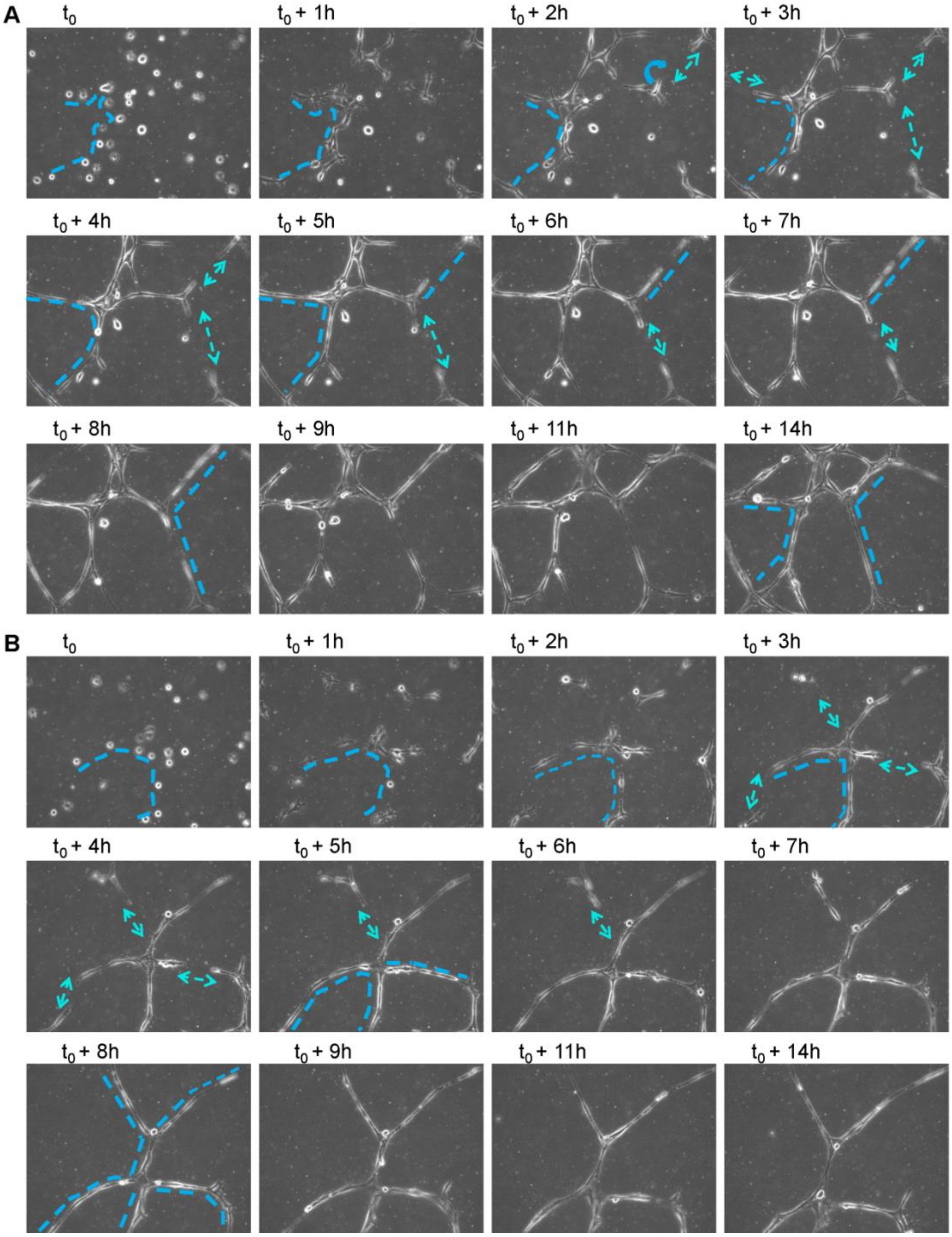


**Figure S2. The dynamic process of ASMCs network assembly on 3D Matrigel containing COL. (A & B)** Two parallel samples are shown. The broken lines indicate the process how the group of cells nearby got connected, formed a curved chain, and further got straightened up. The curved arrows indicate the cell body bending. The double-head arrows indicate that cells could sense each other distantly, and move directionally toward each other, followed by getting connected and straightened up.

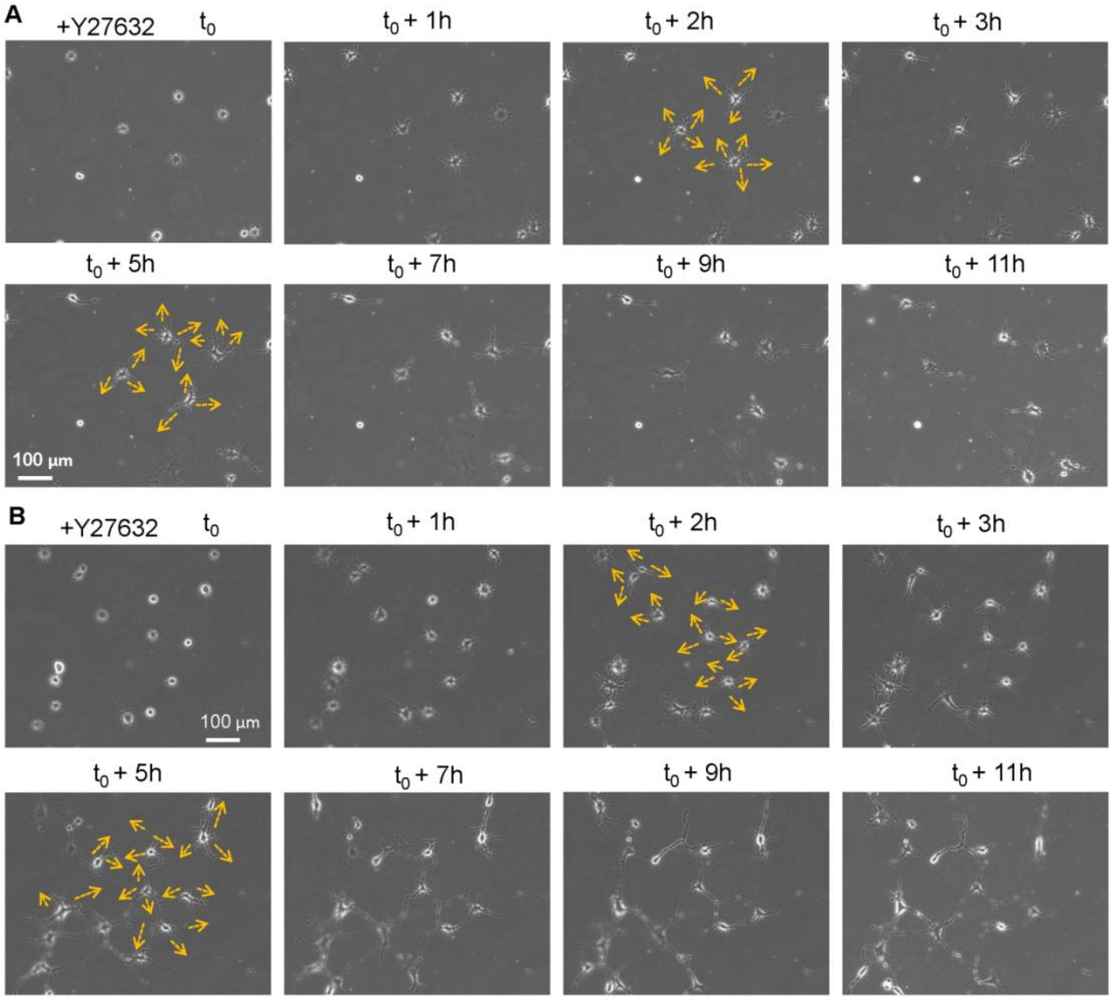


**Figure S3. Inhibition of cell contraction force inhibited directional budding. (A & B)** Cells were seeded on 3D hydrogel containing 0.5 mg/ml COL along with addition of 40 μM Y27632 in the culture medium. (A) and (B) show different local cell densities respectively. The arrows indicate the budding directions not relevant to the positions of other cells nearby.

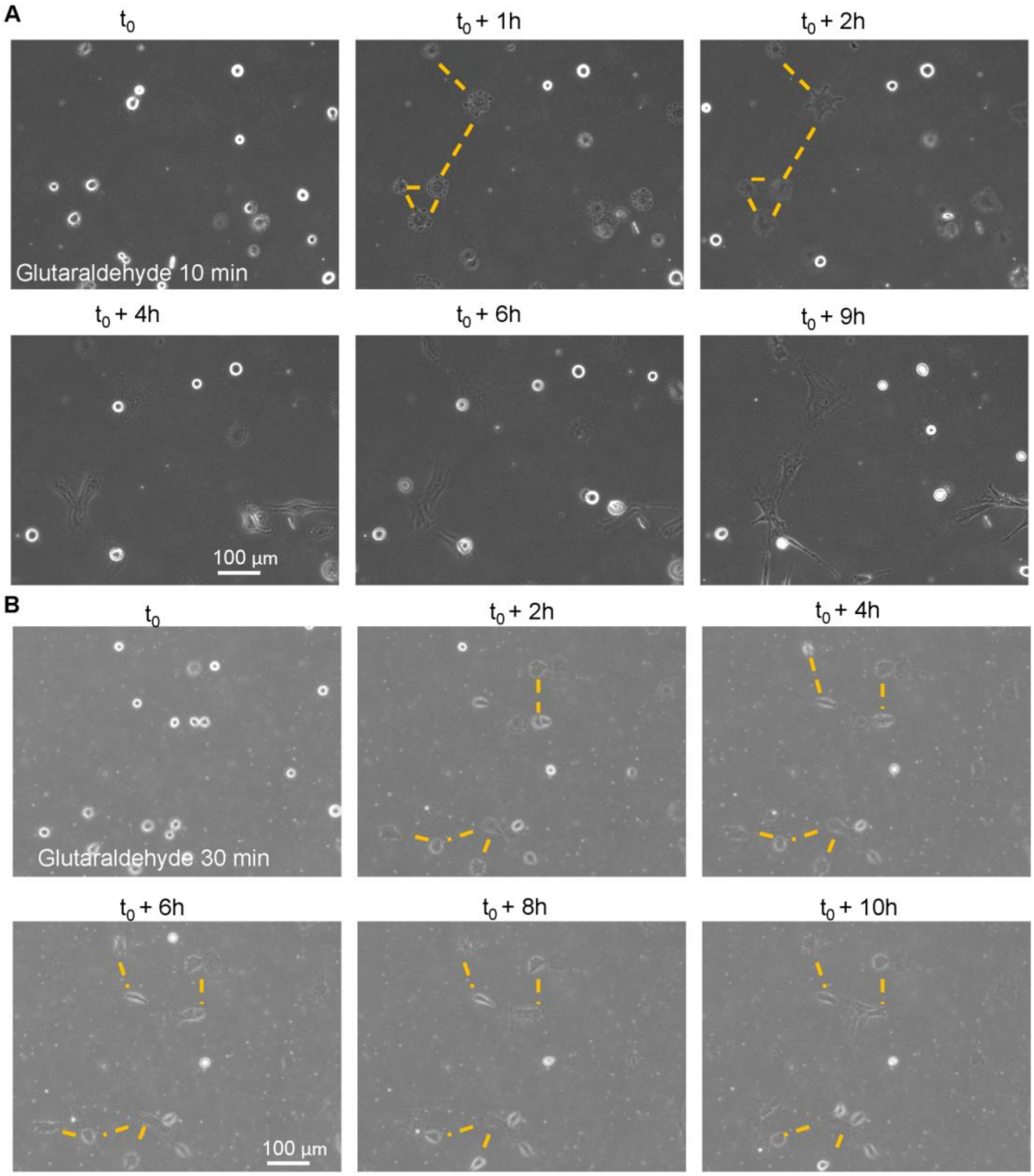


**Figure S4. Cross-linking of the 3D hydrogel inhibited directional budding and cell-cell mutual sensing. (A & B)** The time sequences of cells cultured on 3D hydrogel pre-treated with 0.5% glutaraldehyde solution for 10 min (A) or 30 min (B). The lines indicate the distance between cells.

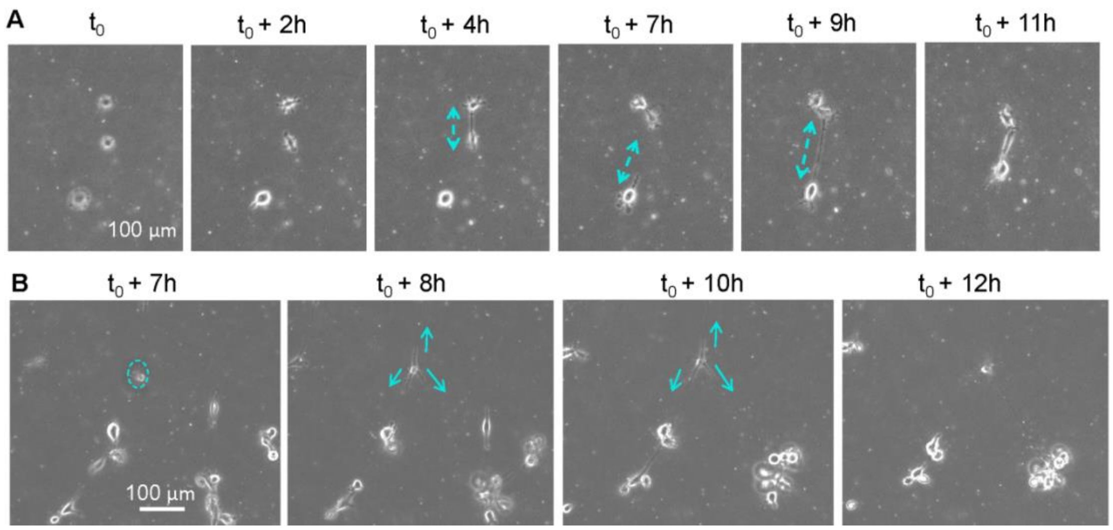


**Figure S5. The distant sensing of ASMCs cultured on non-fibrous 3D Matrigel without COL. (A)** The mutual connection of three isolated cells through slow directional movements. Cells were seeded on 3D Matrigel at low density (~3,000 cells/cm2). **(B)** The cell-cell distant sensing and directional budding, followed by formation of cell clusters at higher cell-seeding density (~9,000 cells/cm2). The arrows indicate cell-cell position sensing and directional movements.

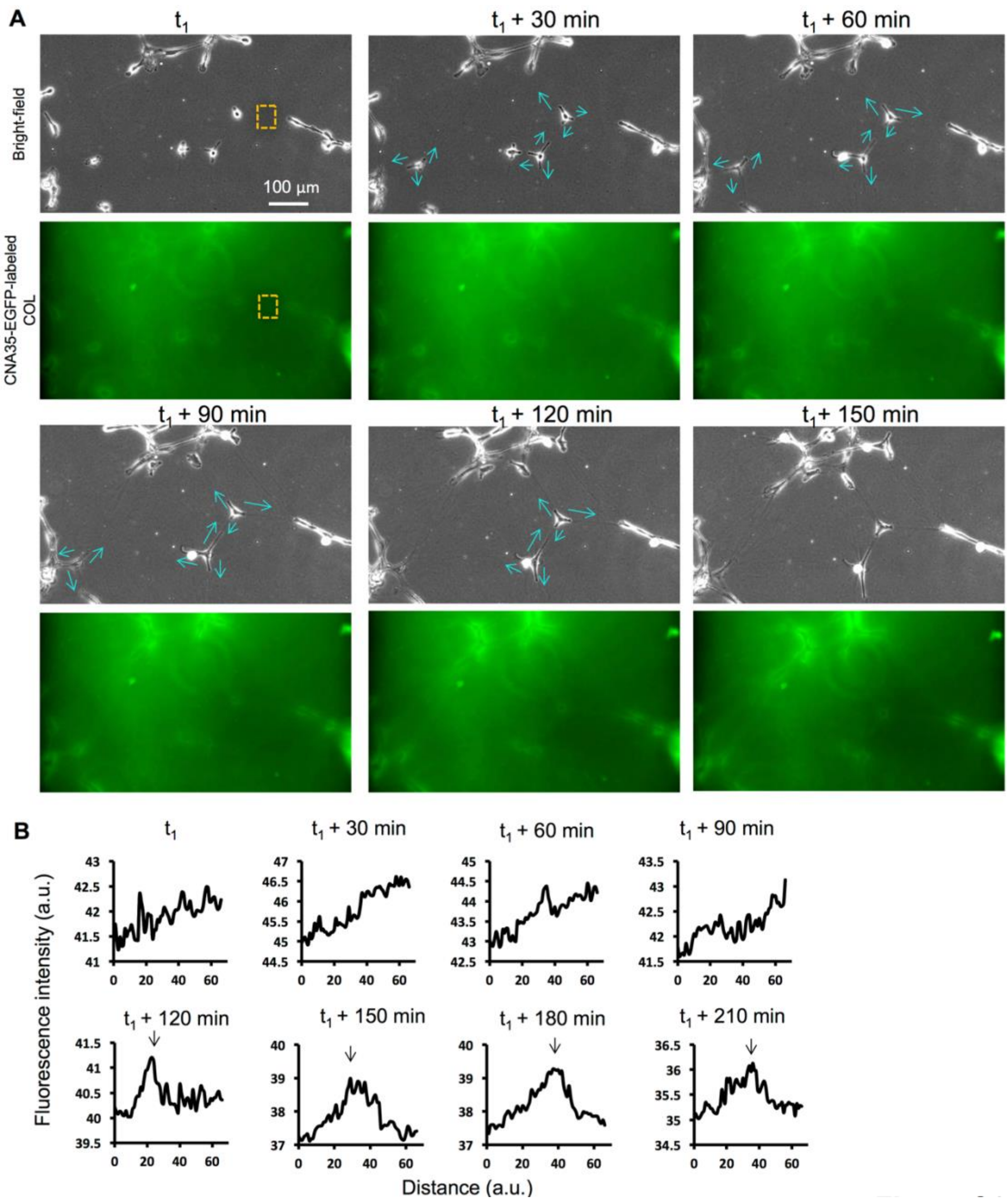


**Figure S6. Cell-cell distant position sensing before COL fiber assembly, and the network formation of COL fibers. (A)** The time-lapse images of three cells which accurately protruded to neighboring cells along with the distribution change of fluorescence-labeled COL. The arrows indicate the directional protrusions. **(B)** The quantifications of fluorescence intensity in the up-downward direction along the selected rectangular region. The arrows point out the intensity peaks on the curves, indicating the gradual accumulation of COL between the two cells with distant sensing.

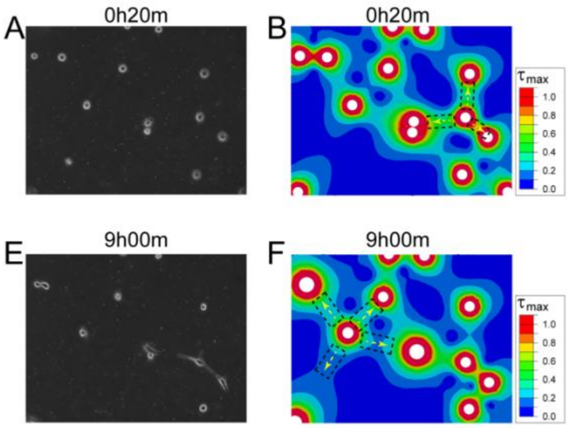


**Figure S7. The directions of cell protrusions were in consistency with the FEM modeling for traction strain distribution. (A)** Cells distribution on matrix at $t_0$+20min, as also displayed on Fig. 4A. **(B)** The distribution of maximum shear stress on the matrix at $t_0$+20min. Three protrusion directions were predicted by maximum shear stress distribution, which were shown as black dash rectangles and yellow dash arrows. **(C,D)** Pseudopodia formed along the prediction directions as shown by yellow dashed arrows at $t_0$+3h and $t_0$+6h. **(E)** Cells distribution on matrix at $t_0$+9h. **(F)** The distribution of maximum shear stress on the matrix at $t_0$+9h. Four protrusion directions were predicted and shown as rectangles and arrows. **(G,H)** Pseudopodia formed along the prediction directions as shown by yellow dashed arrows at $t_0$+16h and $t_0$+18h. The pseudo-color scale bar from blue to red represents increasing maximum shear stress at local place, which indicates high traction strain between cells relatively.

### Supplementary Movie legends

**Movie S1. Distant interactions of ASMCs during network assembly.** The time course shows the dynamic process of ASMCs network assembly on 3D Matrigel containing COL. Interval time = 20 min.

**Movie S2. The directional movements and branching assembly of HUVECs on the hydrogel.** HUVECs were seeded at different densities on 3D Matrigel containing 0.5 mg/ml COL. Time-lapse imaging (interval = 30 min) revealed the dynamics processes of cell-cell distant attractions, cell directional movements, and branching assembly.

**Movie S3. The distant mutual sensing and directional budding of ASMCs.** The time course shows the dynamics process of cell distant mutant sensing and directional budding precisely pointing to neighboring cells in distance during network assembly on 3D Matrigel containing COL. Interval time = 20 min.

**Movie S4. Inhibition of contraction force resulted in cells losing mutual sensing or directional budding.** The time course shows the non-directional budding of ASMCs on 3D Matrigel containing COL when addition of traction force inhibitor (40 μM Y27632). Interval time = 20 min.

**Movie S5. Inhibition of traction force transmission in the matrix led to cells losing mutual sensing or directional budding.** The time course shows losing mutual sensing and directional budding of ASMCs on 3D Matrigel containing COL after the hydrogel was cross-linked by 0.5% glutaraldehyde treatment. Interval time = 30 min.

**Movie S6. No distant mutual sensing or directional budding of ASMCs on coated glass.** The two time courses show no distant mutual sensing or directional budding of ASMCs on COL or Matrigel-coated glass. Interval time = 20 min.

**Movie S7. The distant sensing of ASMCs cultured on non-fibrous 3D Matrigel only.** The time course shows the mutual sensing and directional movements of ASMCs on 3D Matrigel without COL, though it was less efficient than on 3D Matrigel with COL. Interval time = 20 min.
